# Tolerogenic transcriptome landscape in CD8+ T lymphocytes after exposure to erythrocyte-targeted antigens

**DOI:** 10.1101/278457

**Authors:** Alizée J. Grimm, Cédric Gobet, Giacomo Diaceri, Xavier Quaglia-Thermes, Jeffrey A. Hubbell

## Abstract

Our group has recently shown induction of antigen-specific T cell tolerance through targeting of the antigen to erythrocytes *in situ*. The tolerogenic state is characterized by initial proliferation of antigen-specific T cells and subsequent acquisition of signatures associated with both deletional, anergic and regulatory T cell phenotypes. In this study we wished to further understand the molecular mechanisms behind induction of tolerance by erythrocyte-targeted antigens. RNA sequencing was performed to determine how gene expression response is regulated in tolerized ovalbumin-specific CD8+ T cells and which molecular pathways are activated after treatment with this technology. Treatment with erythrocyte-targeted antigens led to the upregulation of genes encoding several TCR co-inhibitory receptors such as CTLA4, PD1, LAG3, TIGIT and CD200R1, and lack of upregulation of cytotoxic and pro-inflammatory signaling molecule genes. Modulation in expression of the master transcription factors *Egr2*/*NFatc1, Nur77* family and *E2f1* was also observed, all known to be associated with the natural process of establishment of peripheral tolerance. Expression of these genes differed in response to treatment with soluble ovalbumin or SIINFEKL MHCI peptide, suggesting a specific mechanism of T cell modulation and tolerance induction in response to the erythrocyte-associated forms.

## Introduction

The development of an antigen-specific therapy is a major step toward treatment of autoimmunity, prevention of graft-versus-host disease or prevention of immune response to protein drug therapeutics. Antigen-specific tolerance strategies have focused on finding means to deliver the antigen to antigen-presenting cells (APCs) in a tolerogenic environment, or in a tolerogenic manner, thus preventing APC maturation and expression of co-stimulatory factors. Antigens associated with self apoptotic cells are known to be processed tolerogenically *in vivo* (1)(2). Based on this property, apoptotic cells have been used by several groups as a carrier of the antigen for induction of immune tolerance (3). Cells harvested from the subjects are rendered apoptotic and coupled to the antigen *ex vivo* and are subsequently re-administered to induce tolerance. Positive outcomes in several animal models of autoimmunity such as type 1 diabetes (4) and multiple sclerosis (5), as well as in allergy (6) and tissue transplantation (7), have pushed the concept toward a first-in-man clinical trial for treatment of multiple sclerosis (8). Administration of autologous peripheral blood mononuclear cells (PBMC) chemically coupled to seven myelin peptides was shown to be safe and tolerated by patients and resulted in a diminution of autoreactive T cells in the blood (8). We sought to utilize this same immunobiology, but in a manner that did not require removal of cells from the subject, *ex vivo* processing and subsequent re-administration into the subject to affect ongoing immunity.

With the goal of exploiting the suppressive potential of apoptotic cells without manipulating cells *ex vivo*, our laboratory has developed a technology that harnesses the erythrocyte as a tolerogenic vehicle. Billions of erythrocytes die every day following a mechanism similar to apoptosis (eryptosis), making them an interesting population to target for induction of immune tolerance (9). To bypass the requirement of *ex vivo* manipulation of the cells, the antigen is engineered such that it will bind to erythrocytes *in situ* after intravenous injection. Working with murine models, intravenous administration of erythrocyte-targeting antigens was shown to lead to T cell tolerance, as measured by to the initial proliferation and subsequent deletion and unresponsiveness of antigen-specific CD4+ and CD8+ T cells (10). The tolerogenic status was maintained over time and dependent on the presence of CD25+ T cells, with upregulation of FOXP3+CD25+ T cells, suggesting a direct role of regulatory T cells (Tregs) in this process (11). In addition to T cell tolerance, prophylactic treatment with erythrocyte-targeted *E. coli* L-asparaginase (which is used to treat acute lymphoblastic leukemia) could prevent the development of a subsequent humoral immune response against this protein therapeutic, a mechanism believed to be due to the antigen-specific deletion of CD4+ T cells and prevention of B cell activation through removal of T cell help (12). The approach of erythrocyte association of antigen has been validated by another approach, that being by enzymatic conjugation of antigens to genetically engineered erythrocytes followed by transfusion (13).

Peripheral activation of self-reactive CD8+ T cells is dependent on the environmental and co-stimulatory signals provided during antigen-presentation (14). Depending on these signals, activation of naïve T cells in the periphery can either result in a functional effector phenotype or T cell tolerance. Tolerance itself regroups several fates such as exhaustion, deletion, anergy, or differentiation into regulatory T cells (15). The molecular signature of T cells undergoing peripheral tolerance has been dissected and the transcriptome responses during exhaustion (16), deletional tolerance (17) or anergy (18) have been characterized. To further understand the molecular mechanisms involved during induction of immunological tolerance by erythrocyte-targeted antigens, the transcriptome status of tolerized CD8+ T cells was assessed by RNA sequencing (RNAseq). Transgenic ovalbumin-specific CD8+ T cells, i.e. OTI T cells, were used in this study and gene response after treatment with erythrocyte-targeted antigens was compared to that in response to equimolar soluble SIINFEKL MHCI peptide, equimolar full-length ovalbumin (OVA) or in a naïve state. To identify genes and pathways critical in the induction of tolerance by erythrocyte-targeted antigens, published datasets of genes upregulated in CD8+ T cells during induction of peripheral tolerance were used and compared to our dataset.

In summary, treatment with erythrocyte-targeted antigens led to the upregulation of genes associated with inhibitory immune functions and exhaustion, and lack of expression of several genes associated with effector functions. In addition, gene response to treatment with erythrocyte-targeted antigens was shown to be comparable to that reported during the natural process of induction of peripheral tolerance, emphasized by the selective modulation of transcription factors associated with TCR signaling and implicated in the process of regulation of apoptosis, such as the *Egr2* and *Nur77* family of transcription factors. Finally, CD8+ T cells harbored the cell cycle-specific features of peripherally tolerized CD8+ T cells, through modulation of the expression of the transcription factor *E2f1* and E2F1 targets.

## Materials and methods

### Animals

All experiments were conducted under the prior approval of the Swiss Veterinary Authority and the EPFL Centre d’Application du Vivant, with all methods being in accordance with the relevant guidelines and regulations. Male C57BL/6 mice (Harlan) aged 8-12 wk were used for *in vivo* studies. CD45.1+ OTI mice were generated by crossing C57BL/6-Tg (TcraTcrb) 1100Mjb (OTI) mice (Jackson Laboratories) with C57BL/6-Ly5.1 (Charles River). Transgenic CD45.1+ OTI mice were bred in specific pathogen-free (SPF) conditions at the Ecole Polytechnique Fédérale de Lausanne Animal Facility, and males were used for splenocyte isolation at 8-12 wk of age.

### ERY1-OVA chemical conjugation

Our laboratory has previously used phage display of peptide libraries to discover a peptide, referred to as ERY1, that binds specifically to murine glycophroin A, which is present only on erythroid cells; this peptide can be chemically conjugated to protein antigens to target them to erythrocytes after i.v. injection (19). 10 equivalents of ERY1 peptide (H_2_N-WMVLPWLPGTLDGGSG-CRG-CONH_2_) were chemically conjugated to ovalbumin (OVA) using a sulfosuccinimidyl-4-(*N*-maleimidomethyl) cyclohexane-1-carboxylate linker (Thermo Scientific). In summary, SMCC was dissolved in dimethylformamide (DMF) and 10 molar equivalents were reacted with endotoxin-free OVA (OVA; Hyglos GmbH) at a concentration of 5 mg/ml in PBS (Dulbecco’s, Sigma-Aldrich) for 2 hr at RT. Unreacted SMCC was removed from the solution using a Zeba desalting column (Thermo Scientific). The OVA-SMCC product was subsequently reacted with 10 molar equivalents of ERY1 peptide dissolved in 3M GndHCl for 2 hr in PBS. The final product was loaded in a Zeba desalting column to remove unreacted ERY1, sterile-filtered and stored at -20°C.

### TER119-SIINFEKL recombinant protein expression

Our laboratory has previously reported use of a murine glycophorin A-binding antibody, referred to as TER119, to target recombinantly fused antigens to erythrocytes after i.v. injection (10). The TER119 scFv antibody fragment was created as described elsewhere (10) and inserted in a PsectagA expression vector (Life Technologies). TER119-SIINFEKL (comprising the 7 N-flanking amino acids from OVA to allow for antigen processing) was expressed in suspension culture of HEK293E cells under serum-free conditions with 3.75 mM valproic acid (Sigma-Aldrich) for 7 d. Proteins were purified from supernatant using immobilized metal ion affinity chromatography on an Akta FPLC system (GE Healthcare) and by size-exclusion using a Superdex 75 column (GE Healthcare). After purification, purity was assessed by SDS/PAGE, concentration was determined by nanodrop and endotoxin levels were measured using THP-1 × Blue cells (InvivoGen). Final products were sterile-filtered and stored at -80 °C.

### Mouse study

CD8+ T cells from the spleen of OTI mice were isolated using a CD8+ magnetic bead negative selection kit (Miltenyi Biotec), following the manufacturer’s instructions. On day 0, 10^6^ CFSE-labeled CD8+ OTI cells were injected i.v. in 100 μL of PBS into the tail vein of recipient CD45.2+ C57BL/6 mice. One day following adoptive transfer, 223 pmol (equivalent to 10 μg of OVA) of ERY1-OVA, TER119-SIINFEKL, OVA or SIINFEKL peptide were injected i.v. in a 100 μL volume of saline. Mice were euthanized on day 4 and spleens were harvested for OTI CD8+ T cell isolation. To obtain a sufficient quantity of RNA for sequencing, spleens from 3 mice in the same treated group were pooled before cell sorting. From 9 mice per treatment group, we obtained 3 samples per group to be sent for RNA sequencing.

### Cell sorting and RNA isolation

Splenocytes were exposed for 5 min at RT to 0.155 M NH_4_Cl to lyse erythrocytes. A first round of isolation was performed for CD8+ T cell isolation using a CD8+ magnetic bead negative selection kit (Miltenyi Biotec), following the manufacturer’s instructions. The following staining steps were performed on ice. Cells were washed with PBS and stained for 15 min in PBS + 2% FBS with CD3e FITC, CD45.1 PE-Cy7, CD8 APC-e780, and CD4 Pacific blue antibodies (eBioscience). Cells were resuspended at a concentration of 8x10^6^ cells/mL in PBS + 1 mM EDTA, and OTI CD8+ T cells were sorted using a FACS ARIAII cytometer. Viable transgenic OTI CD8+ T cells were gated based on their expression of the CD3e, CD8 and CD45.1 markers (Fig. S1). Total RNA was extracted from sorted OTI CD8+ T cells using a RNeasy Plus Micro kit (Qiagen), following the manufacturer’s instructions.

An average of 21,000-52,000 OTI T cells were sorted per sample. The quality and quantity of extracted RNA was analyzed by a fragment analyzer (Fig. S2A). RNA integrity was very satisfying, except for sample SIIN 1, from the SIINFEKL-treated group, which was partially degraded. Subsequent generation of the library failed for this sample, leaving n=2 for the SIINFEKL group in this study.

### RNA sequencing

Quantification and qualification of extracted RNA was assessed by capillary electrophoresis using a fragment analyzer. Libraries were generated using an Illumina RNA-SEQ kit (TruSeq^®^ Stranded mRNA LT – SetA, RS-122-2101) and sequencing was performed on an Illumina HiSEQ4000 for a single-end 50 cycles run. Demultiplexing was done using Illumina “bcl2fastq2-v2.17.1.14” software.

### RNA-Seq mapping and quantification

Single end reads were aligned to the mouse genome (assembly GRCm38/mm10) using STAR 2.4.2.a (18) with default parameters. A genome index was built using Ensembl annotation to improve splice junction accuracy. For each Ensembl gene, all of the exons of the respective annotated protein-coding transcripts were considered. Using a custom Perl script, reads up to one mismatch were counted, considering only reads in the right gene orientation. Both uniquely mapping and multimapping reads were reported. Genes were not considered for analysis if the average number of read was lower than 100 reads in more than two conditions, or in the case of genes with less than 10 reads in a certain condition. Read counts were normalized by the respective library size using EdgeR (19), and by the average size of transcripts (RPKM). Log2 gene expression levels were calculated and reported.

### Model selection and evaluation of differential gene expression

In the interest of clustering genes exhibiting similar behavior in the five conditions, i.e. naïve, ERY1-OVA, OVA, TER119-SIINFEKL and SIINFEKL, we used a model selection approach under the generalized linear model framework of EdgeR. For this purpose, all possible additive combinations of factors and their respective design matrices were generated, leading to a set of 52 different modules (Fig. S3). Each module was solved using generalized linear regression. The Aikake information criterion (AIC) was computed via the deviance of the resulting fit. Finally, all the genes were assigned to their preferred module based on their minimal AIC value. For simplification and enrichment analysis, modules harboring similar responses were grouped together, leading to a total of 16 models clustering the genes specifically expressed in response to the different treatments. Moreover, genes were split in accordance to a positive or negative regulation compared to the naïve group. For the sake of comprehension, the different models are named based on the first letters of the treatments inducing expression of the genes clustered in the model compared to naive, i.e. E for ERY1-OVA, O for OVA, T for TER119-SIINFEKl and S for SIINFEKL, followed by the model number. For instance, EOT_13 model regroups genes of which expression is induced in response to ERY1-OVA, OVA and TER119-SIINFEKL, but not in response to treatment with SIINFEKL peptide. Log2 normalized read counts (RPKM) of genes and attributed models can be found in the supplementary file 1.

### Gene ontology (GO) term analysis

Gene ontology analysis was performed using the TopGO R package (Alexa *et al*., 2016). Enrichment analysis for GO terms derived from "Biological Process" ontology was done in the different models for upregulated and downregulated genes and significance assessed using the Fisher exact test. GO terms with p-value < 0.01, a minimum number of 2 genes and less than 750 annotated genes were considered. Genes with more than 10 reads on average across all conditions were selected as background for the analysis. All GO terms derived from the gene ontology analysis can be found in the supplementary file 2 and 3 for up- and down-regulated respectively.

## Results

### Cell sorting and RNA extraction

Regulation of gene expression in CD8+ T cells after treatment with soluble versus erythrocyte-targeted antigens was assessed using as a T cell model OTI T cells, i.e. CD8+ T cells bearing a transgenic T cell receptor specific for the SIINFEKL sequence in OVA. In summary, 10^6^ naïve CFSE-labeled OTI T cells were adoptively transferred in C57BL6/J mice on day 0. One day following adoptive transfer, mice were treated by intravenous administration of soluble MHCI SIINFEKL peptide, full-length OVA, or their erythrocyte-targeted forms, i.e. TER119-SIINFEKL and ERY1-OVA. Spleens were harvested on day 4, and OTI CD8+ T cells were isolated for RNA extraction (Fig. 1A). Live OTI T cells were selected based on their expression of the surface markers CD3e, CD8 and CD45.1 (Fig. S1). We selected a time-point of 4 days post-treatment, as we wished to observe the maximal effect of the different treatments on gene modulation in CD8+ T cells while avoiding substantial cell death. Indeed, CD4+ and CD8+ T cells tolerized by erythrocyte-targeted antigens undergo an initial proliferative phase followed by cell death or unresponsiveness that can be measured as soon as 5 days post-treatment. Significant cell death would impact the quality of the results obtained by RNA sequencing and should thus be avoided.

**Figure 1:**
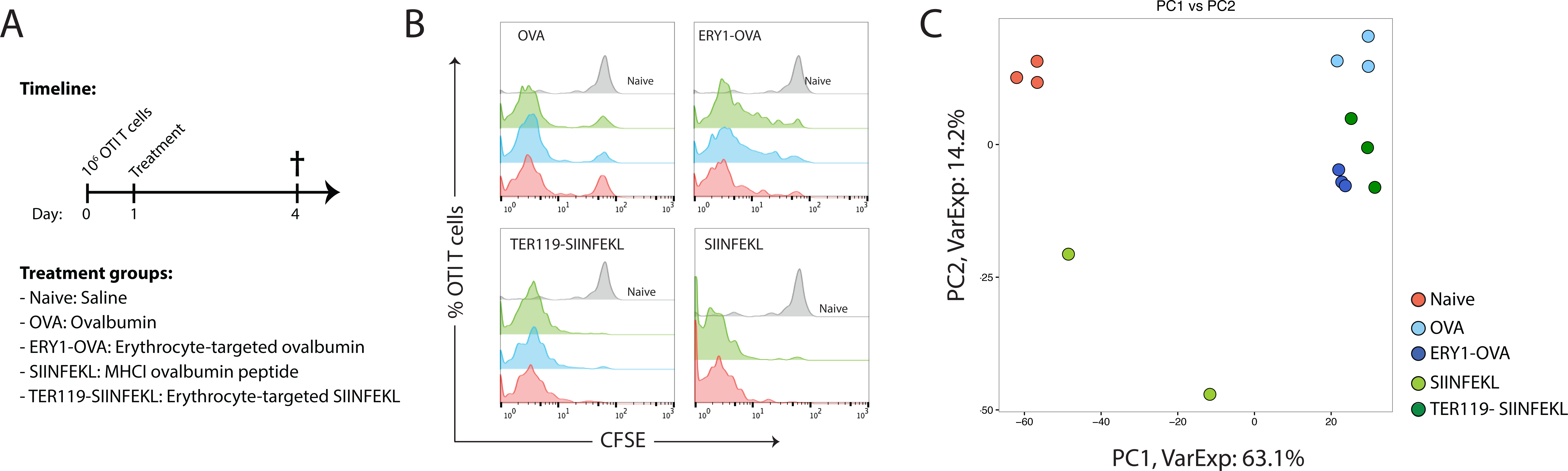
Distinct gene response in CD8+ T cells after treatment with erythrocyte-targeted or soluble antigens. (A) 10^6^ naïve OTI T cells were adoptively transferred in C57BL6/J mice on day 0. One day following adoptive transfer, mice were treated by intravenous administration of soluble MHCI SIINFEKL peptide, full-length OVA, or their erythrocyte-targeted forms, i.e. TER119-SIINFEKL or ERY1-OVA. Spleens were harvested on day 4, and OTI CD8+ T cells were isolated for RNA extraction and sequencing. (B) Flow cytometry histogram representation of CFSE dilution in proliferated OTI T cells on day 4. (C) Principal component analysis: each dot represents one of the 14 samples in a multidimensional gene expression space projected on the first and second principal component. A total of 77.3% of the variance of the samples was comprised within the first two components.

### Distinct gene regulation in response to treatment with erythrocyte-targeted versus soluble antigens

RNA extracted from CD8+ T cells in mice treated with soluble or erythrocyte-targeted antigens was analyzed by RNA sequencing (RNAseq), and compared to that of naïve resting T cells in mice treated with saline. As previously reported (10), administration of all the different treatments induced proliferation of OTI T cells, as measured by CFSE dilution on day 4 (Fig.1B). Samples from the same treatment groups co-localized in the two first dimensions of the principal component analysis (PCA), suggesting consistency in the gene response within each treated group, with highest variability in the SIINFEKL group (Fig. 1C). PCA revealed that CD8+ T cells tolerized by erythrocyte-targeted antigens, soluble OVA or MHCI peptide SIINFEKL, harbored a distinct gene signature from that of naïve CD8+ T cells, but also between each other (Fig. 1C). High proximity in the PCA for the ERY1-OVA and TER119-SIINFEKL conditions was observed, suggesting a comparable modulation of CD8+ T cells after treatment with the two erythrocyte-targeting strategies, i.e. chemical conjugation with ERY1 peptide or recombinant expression of a TER119 scFv fusion protein. The first component of the PCA was shown to segregate the naïve group from the treated groups, while the second component separates the SIINFEKL treated group from the others (Fig. 1C). Replicates from the same condition exhibited high reproducibility, as measured by correlation analysis (Pearson coefficient 0.924-0.99) (Fig. S2B).

Model selection was performed, and out of the 9931 genes expressed at a quantifiable level, a total of 6056 genes were identified as being differentially expressed in response to at least one of the conditions compared to naïve (Fig. S2C). Genes were clustered into 16 distinct models based on their relative level of expression in response to the different treatments, and were finally split in accordance to a positive or negative regulation compared to the naïve group (Fig. 2A-B). To provide information regarding the biological processes induced after administration of erythrocyte-targeted or soluble antigens, gene ontology analysis was performed on the different models (Fig. 2C). For genes for which expression was downregulated compared to naïve, heatmap representation and results from the gene ontology analysis can be found in the supplementary figure S4.

**Figure 2:**
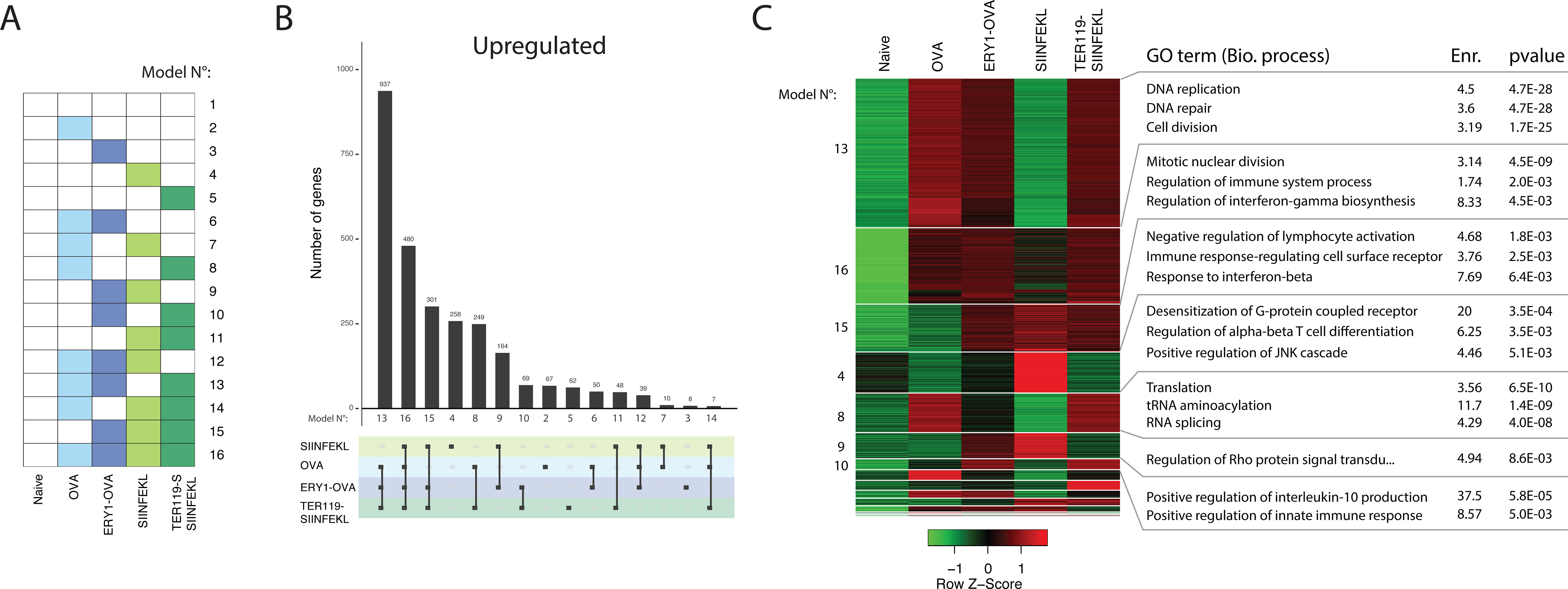
Model selection and gene ontology analysis were performed to segregate genes exhibiting similar behavior. (A) Genes were classified into 16 models, based on their level of expression in response to the different treatments. Colored boxes represent genes expressed differentially compared to the naïve group. (B) Top: number of genes in each model is represented as a bar plot for genes upregulated relatively to the naïve group. Bottom: combination matrix encoding the respective conditions and assigned model number. (C) Heatmap illustrating the z-score of genes upregulated relative to the naïve group. High versus low relative level of expression are depicted in red and green, respectively and assigned model number is shown on the left of the heatmap. Gene ontology analysis and selected GO terms are depicted on the right. For Naïve, ERY1-OVA, OVA and TER119-SIINFEKL, n=3; for SIINFEKL, n=2.

Gene ontology analysis revealed that all treatments, i.e. both erythrocyte-targeted and soluble antigens, induced expression of genes associated with cell cycle and immune regulation (model EOTS_16, Fig. 2C). This enrichment is in concordance with the highly proliferative state of CD8+ T cells measured by flow cytometry under all treatment conditions (Fig. 1C).

A significant fraction of the genes, i.e about 11%, were shown to be specifically upregulated after treatment with erythrocyte-targeted antigens and soluble OVA, but not in response to soluble SIINFEKL peptide (model EOT_13, Fig. 2B-C). This observation matches the very distinct gene signature of the SIINFEKL group observed from the first two components of the PCA (Fig. 1C). GO analysis of cluster EOT_13 highlighted strong enrichment of cell-cycle genes (Fig. 2C) as well as downregulation of genes associated with Ras pathway, known to be associated with TCR signaling and T cell activation (20)(21) (Fig. S4).

The third most represented model was shown to be model ETS_15, i.e. genes upregulated by all conditions beside OVA. Thus, a significant fraction of genes were specifically upregulated by erythrocyte-targeted but not soluble OVA and involved in the process of immune system response (GO analysis model ETS_15, Fig. 2C).

Finally, gene ontology analysis revealed an interesting behavior in the set of genes for which expression was specifically induced in response to erythrocyte-targeted antigens, i.e. ERY1-OVA and TER119-SIINFEKL, but not in response to soluble OVA or SIINFEKL peptide (model ET_10). Indeed, model ET_10 depicted strong enrichment for genes associated with the positive regulation of IL-10 production, a key anti-inflammatory cytokine expressed by regulatory T cells (Fig 2C).

### Molecular parallels between anergy, deletional tolerance, exhaustion and response of CD8+ T cells to erythrocyte-targeted antigens

The transcriptome programs of activated and tolerized CD8+ T cells have been investigated by others, and groups of genes associated with these events have been reported. To understand and dissect the response induced after treatment with soluble or erythrocyte-targeted antigens, our dataset was compared to gene sets characteristic of activated and tolerized T cells. Genes specifically upregulated in memory (clusters 2 & 10, Schietinger *et al*.(18)) or rescued/effector CD8+ T cells (clusters 2 & 7, Schietinger *et al.*(18)) were used as signature for T cell activation. In parallel, genes specifically upregulated during deletional tolerance (17), peripheral self-tolerance/anergy (clusters 9 & 13, Schietinger *et al.*(18)), exhaustion in chronic viral infection (16) or in exhausted tumor-specific T cells (22) were used to evaluate the subtype of tolerogenic response induced by the different conditions. Genes upregulated in response to the different treatments were tested for overrepresentation in these published dataset. Gene enrichment analysis was performed to *i*) determine if treatments with erythrocyte-targeted or soluble antigens induced expression of genes reported as upregulated during immune activation or tolerance (Fig. 3A), and *ii*) understand the combined responses using our model classification (Fig. 3B).

**Figure 3:**
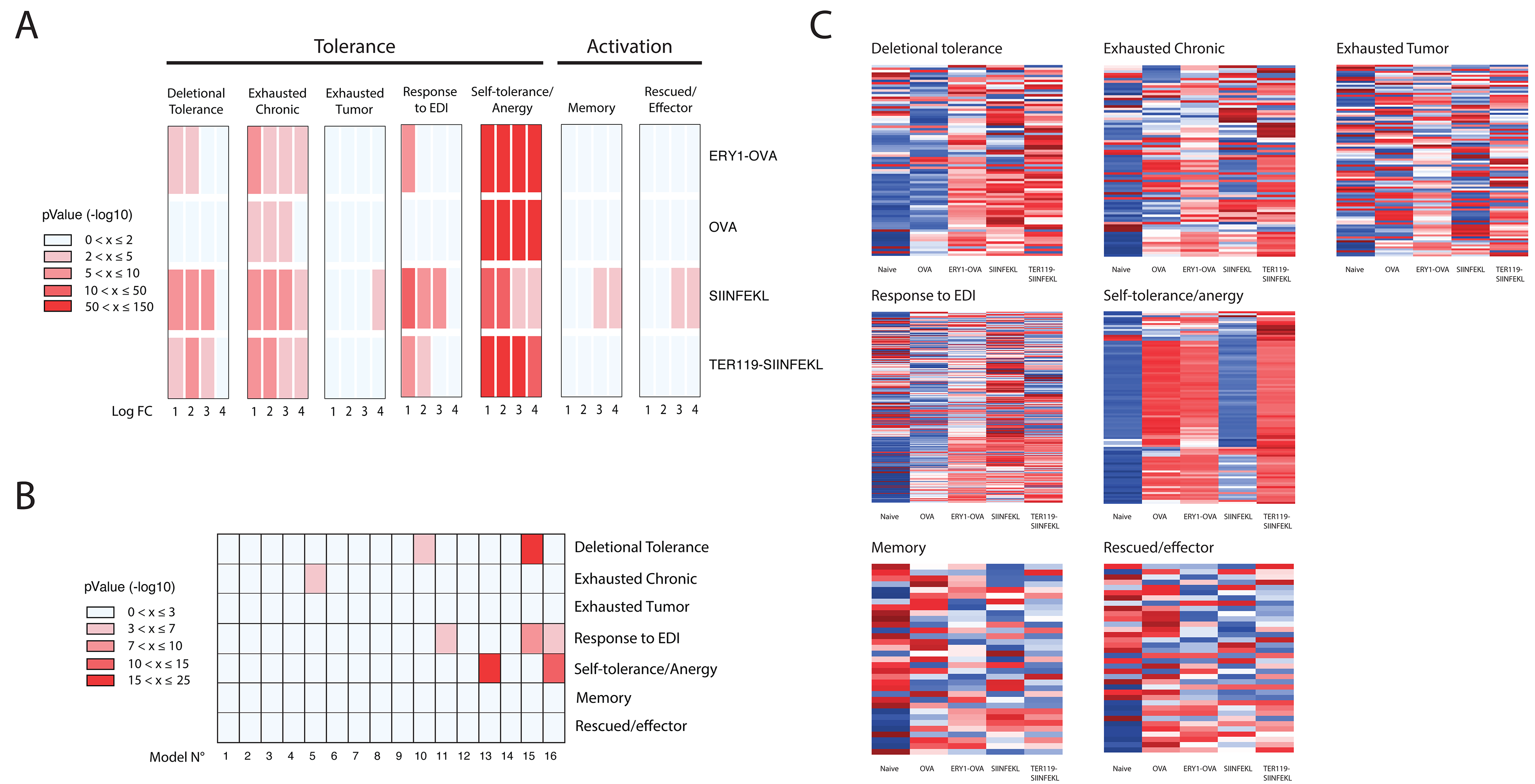
Specific tolerogenic signature induced after administration of erythrocyte-targeted or soluble antigens. Genes known to be upregulated in memory (clusters 2 and 10 Schietinger et al.(18)) or rescued/effector CD8+ T cells (clusters 2 and 7 from Schietinger et al.(18)) were used as a signature for T cell activation. In parallel, and to represent the different types of tolerogenic responses, genes specifically upregulated during deletional tolerance (17), peripheral self-tolerance/anergy (clusters 9 and 13 from Schietinger et al.(18)), exhaustion in chronic viral infection(16) (abbreviated exhausted chronic in the figures), exhausted tumor specific T cells (22) (abbreviated exhausted tumor) or after escalating dose immunotherapy with soluble peptide (23) (abbreviated response to EDI) were used. Gene enrichment analysis was performed to determine if (A) treatments with erythrocyte-targeted or soluble antigens induced gene regulation patterns fitting a specific activation or tolerogenic response, as well as (B) validate the specific response to each condition through model selection. (C) Relative expression (Z-score) of genes reported as being upregulated during the different activation or tolerogenic responses. High versus low relative level of expression are depicted in red and blue, respectively.

None of the treatments, i.e. ERY1-OVA, OVA, TER119-SIINFEKL or SIINFEKL, induced upregulation of the group of genes reported during T cell activation, i.e. genes upregulated in memory CD8+ T cells or rescued/effector CD8+ T cells (Fig. 3A-B). Instead, overexpression of genes associated to a tolerogenic fate was observed after administration of both soluble and erythrocyte-targeted antigens (Fig. 3A).

Treatment with erythrocyte-targeted antigens led to the enrichment of genes reported to be overexpressed during deletional tolerance, self-tolerance/anergy as well as exhaustion during chronic viral infection (Fig. 3A). Such specific modulation can be appreciated from the heatmap representations of the different comparisons (Fig. 3C). The similarity in gene modulation reported during self-tolerance/anergy by Schietinger and his group (18) and the one observed in our study in response to treatment with erythrocyte-targeted antigens and soluble OVA was remarkable. Indeed, genes associated with self-tolerance were highly overrepresented in response to ERY1-OVA, TER119-SIINFEKL and OVA (Fig. 3A). Interestingly, treatment with SIINFEKL peptide also induced modulation of these genes, but with a much lower amplitude (Fig. 3C). Consistent with this observation, high enrichment in model EOT_13, i.e. genes induced by all treatments beside SIINFEKL peptide, was observed for this subset of genes (Fig. 3B). Together, these analyses revealed that treatment with erythrocyte-targeted antigens and OVA lead to a gene response that is highly similar as the one described during induction of self-tolerance.

While administration of OVA induced expression of genes associated with both self-tolerance/anergy and exhaustion (Fig. 3A-C), no significant overrepresentation of genes reported as being upregulated during deletional tolerance was observed in this group (Fig. 3A). This specific response was supported by enrichment in model ETS_15 for the “deletional tolerance” set of genes (p-value: 2.82E-16) (Fig. 3B).

Finally, and in order to compare erythrocyte-targeting strategy to an antigen-specific immunotherapy currently being evaluated in clinic, our dataset was compared to the transcriptome signature of T cells tolerized by escalating dose immunotherapy (EDI) (23). Through administration of increasing doses of the peptide antigen, EDI has been shown to induce antigen-specific T cell tolerance and regulatory T cells (24). This therapy is currently used in clinic for allergen desensitization and its potential for treatment of autoimmune diseases is also being evaluated (23). Genes significantly upregulated after treatment with erythrocyte-targeted antigens and SIINFEKL peptide, but not soluble OVA, were shown to be overrepresented in the set of genes upregulated after EDI (Fig. 3A). This behavior was emphasized by a significant enrichment in model ETS_15 for this set of genes (p-value: 1.25E-9) (Fig. 3B).

In summary, through comparison of our dataset with the gene sets characteristic of T cells undergoing either activation or tolerance, treatment with erythrocyte-targeted antigens was shown to induce the modulation of genes associated with tolerogenic responses of both deletional tolerance, exhaustion and self-tolerance/anergy. This signature, and thus type of tolerogenic-response induced, was shown to differ in response to soluble SIINFEKL peptide and OVA. Deeper analysis of the genes, pathways and transcription factors involved in these processes are discussed in detail below.

### Modulation of inhibitory receptors and pro-inflammatory cytokines genes

Functional impairment in self-reactive T cells involves *i*) expression of genes associated with inhibitory immune function or exhaustion, *ii*) the inability to express genes associated with effector functions and *iii*) an altered expression of master transcription factors (18). Fig. 4A offers a summary of salient changes in gene expression associated with the erythrocyte-targeted antigens, and those genes underlined were uniquely regulated by erythrocyte-targeted versus free antigens.

**Figure 4:**
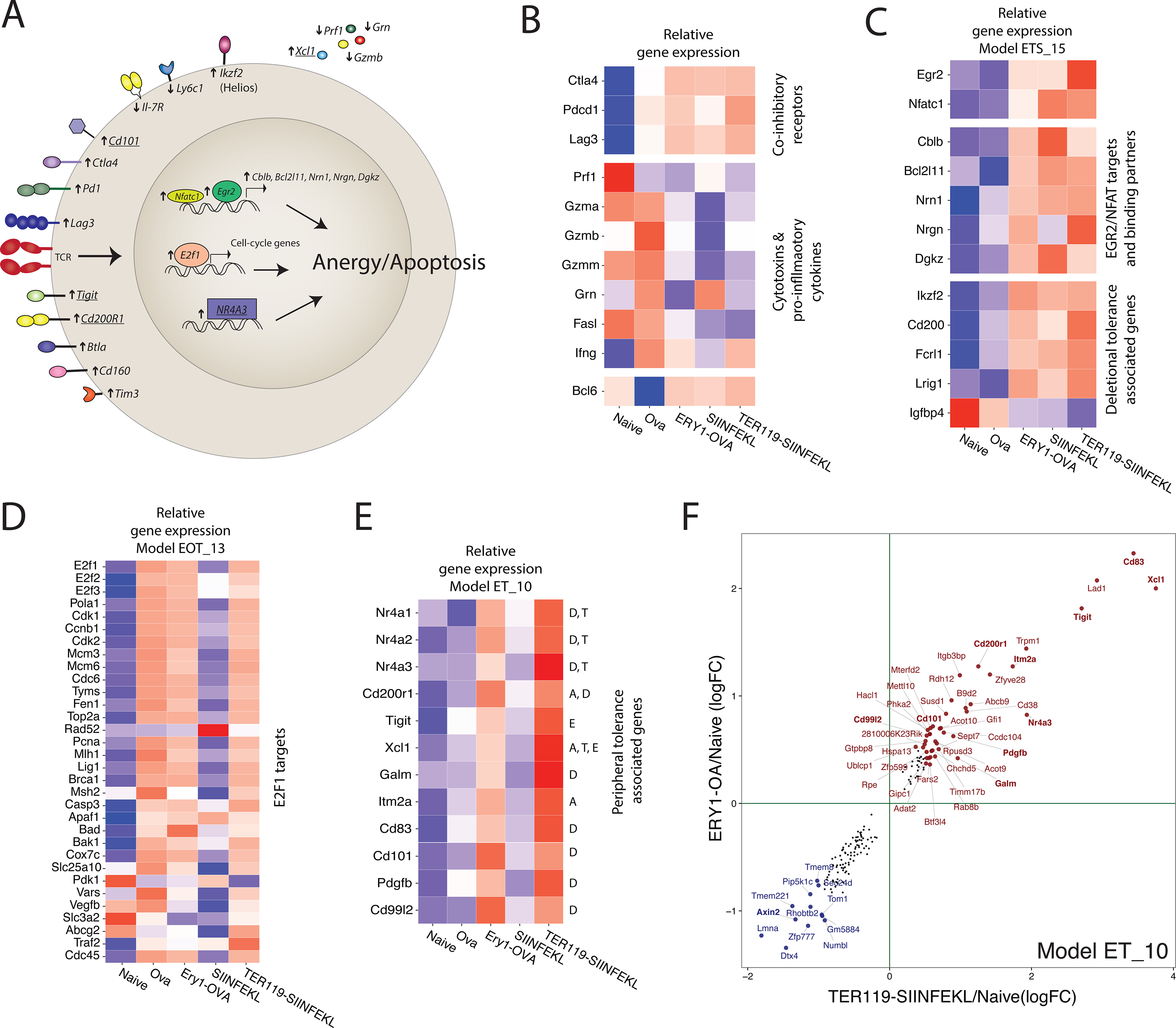
Regulation of inhibitory receptors, cytotoxic molecules genes and master transcription factors in response to erythrocyte-targeted, soluble OVA or SIINFEKL MHCI peptide. (A) Summary of genes and pathways involved in the process of tolerance induction by erythrocyte-targeted antigens. (B) Heatmap depicting the relative gene expression (Z-Score) in CD8+ T cells in response to the different treatments for genes encoding for co-inhibitory receptors and cytotoxic and pro-inflammatory cytokines. (C) Erythrocyte-targeted antigens and soluble MHC I peptide induce regulation of genes associated with deletional tolerance and EGR2/NFATc1 pathway. Relative expression (Z-score) of genes up/downregulated by all treatments beside OVA (model ETS_15). (D) Relative gene expression level (Z-Score) of E2f1 and E2f1 targets (40). (E) Relative level of expression (Z-Score) of genes associated to peripheral tolerance and more precisely to: “A”: anergy (18)(34)(42), “D”: deletional tolerance (17), “E”: Exhaustion (43)(22)(44) or “T”: Treg induction (45)(46). (F) Scatterplot representation of genes expressed specifically in response to erythrocyte targeted antigens but not soluble OVA or SIINFEKL peptide (model ET_10). Genes are depicted with either a red or blue dot, depending if they are up- or down-regulated, respectively, compared to naïve (names of genes with a log2 fold change above +0.5 or below -1 are depicted). For heatmap representation, high versus low levels of gene expression are depicted in red and blue, respectively.

When looking at expression of genes encoding for inhibitory receptors, all treatments were shown to induce expression of the co-inhibitory receptor genes cytotoxic T lymphocyte antigen-4 (*Ctla4*), programmed cell death protein 1 (*Pdcd1, Pd1*) and lymphocyte-activation gene 3 (*Lag3*) (Fig. 4B). While expression of these genes was observed in response to all treatments, upregulation of the co-inhibitory receptor *Cd200r1* (25) and immunoglobulin (Ig) inhibitory receptor *Tigit* (26) were solely induced in response to erythrocyte-targeted antigens (Fig. 4E).

Upon recognition of their target, cytotoxic CD8+ T cells release cytotoxins to induce death of the infected or damaged cell. One of the hallmarks of CD8+ T cells undergoing immune tolerance is their inability to produce pro-inflammatory cytokines and cytotoxins upon antigen-restimulation. Using murine models, we have shown that T cell tolerized by erythrocyte-targeted antigens have a reduced IFNγ, TNFα and granzyme B production upon *in vitro* antigen restimulation (10, 11). When looking at gene expression levels, CD8+ T cells treated with erythrocyte-targeted antigens failed to overexpress genes encoding for perforin 1 (*Prf1*), granzymes (*Gzma, Gzmb* and *Gzmm)*, granulin (*Grn*) as well as Fas ligand (*Fasl*) (Fig. 4B). Surprisingly, while reduced secretion of IFNγ upon antigen-restimulation has been reported after treatment with erythrocyte-targeted antigens (10), no significant change in expression of *Ifng* was observed at the gene level (Fig. 4B). This discrepancy between mRNA accumulation levels versus protein expression of IFNγ during tolerance induction has been reported by others, and blockade of IFNγ production during tolerance was suggested to be mediated post-transcriptionally (17)(16).

Interestingly, CD8+ T cells in mice treated with soluble OVA showed increased expression of *GzmB*, which was consistent with what was previously observed at the protein level (11) (Fig. 4B). This increase in expression did not correlate with an increase in expression of transcription factors related to granzyme expression or cytotoxic effector functions of CD8+ T cells, such as Eomes or Notch2 (27). However, upregulation of *GzmB* could be explained by the significant decrease in expression of the granzyme repressor BCL-6 (28) (*Bcl6*) observed in response to treatment with OVA (Fig. 4B). In addition, AKT1 (*Akt1*) and T-bet (*TBX21*), both involved in the expression of IFNg (29)(30), were shown to be solely upregulated in response to OVA.

### Deletional tolerance signature: involvement of EGR2/NFATc1 pathways

Most of the genes described as being upregulated during deletional tolerance were also shown to be upregulated in response to soluble MHCI peptide and erythrocyte-targeted antigens, but not in response to soluble OVA (Fig. 3). To further dissect the transcriptome response and understand which pathways related to immune tolerance are activated by all treatments beside OVA, genes fitting model ETS_15 were analyzed. Discrimination between T cell activation and induction of T cell tolerance involves over-activation of calcium/nuclear factor of activated T cells (NFAT) signaling and defective capacity to signal through the Ras/ERK pathway upon TCR activation (20)(21). Genes in model ETS_15 were examined for any potential transcription factor associated with the TCR signaling pathway, and genes encoding for the transcription factors early growth response 2 (*Egr2*) and nuclear factor of activated T cells (*NFATc1*) were shown to cluster in this model and be specifically upregulated by all treatments beside OVA (Fig. 4C). TCR engagement leads to the activation of the NFAT pathway and subsequent regulation of the transcription of EGR transcription factor family (31), where NFATc1 complexes with EGR2 and plays together a decisive role in modulating T cell response toward tolerance (32). An increase in *Egr2* expression correlated with the upregulation of its targets E3 ubiquitin ligase *Cblb* (33), pro-apoptotic molecule BIM (*Bcl2l11*) (34)(35), Neuritin 1 (*Nrn1*) (34), Neurogranin (*Nrgn*) (34), as well as EGR2-binding partner DGKZ (*Dgkz)* (36) (Fig. 4C).

Other genes, not directly related to Ras/ERK or EGR2/NFATc1 signaling pathways, but described as being specifically regulated during deletional tolerance (17), followed a similar pattern of expression. Among these, upregulation of the transcription factor Helios (*Ikzf2)*, the immunosuppressive molecule *Cd200*, Fc receptor-like 1 (*Fcrl1)*, Leucine-rich repeats and immunoglobulin-like domains 1 (*Lrig1*) as well as downregulation of insulin-like growth factor binding protein 4 (*Igfbp4*) were observed in response to all treatments besides OVA (Fig. 4C).

In summary, treatment with both erythrocyte-targeted antigens and soluble SIINFEKL peptide induced a similar response in CD8+ T cells as the one observed during deletional tolerance, with notable upregulation of genes associated to the EGR2-NFATc1 pathways. This deletional signature was not induced in response to treatment with soluble OVA.

### Regulation of cell-cycle program through E2F1 deregulation

Peripherally tolerized CD8+ T cells maintained in an anergic state exhibit a specific cell-cycle program, with regulation of genes encoding for proteins involved in mitosis and nuclear division (17)(18). Comparison of our dataset with the list of genes reported as being expressed in self-reactive CD8+ T cells maintained in an unresponsive state in the periphery, i.e. genes in cluster 9 and 13 from Schietinger *et al*. (18), revealed that both treatment with erythrocyte-targeted antigens and soluble OVA, but not soluble SIINFEKL peptide, induced a similar response in CD8+ T cells (Fig. 3). Indeed, most of the genes reported as being upregulated in tolerized CD8+ T cells were shown to be upregulated by treatment with both erythrocyte-targeted antigens and soluble OVA, but to a much lower amplitude in response to soluble SIINFEKL peptide at equimolar dose. This trend was confirmed by enrichment analysis (hypergeometric test), where overrepresentation of tolerance-specific genes was observed in model EOT_13 (Fig. 3B).

Gene ontology analysis of model EOT_13 revealed enrichment in genes involved in cell-cycle processes such as cell division or DNA replication (Fig. 3C), and overexpression of the E2F1 transcription factor was observed in response to all treatments besides SIINFEKL (Fig. 4D). E2F1 transcription factor plays a predominant role in the orchestration of cell-cycle through regulation of the entry in the S-phase and DNA replication (37). In addition, deregulation of E2F1 expression is known to affect induction of apoptosis during peripheral tolerance (18)(38)(39). Treatment with OVA and erythrocyte-targeted antigens induced expression of *E2f1* (Fig. 4D). To support the implication of the E2F1 transcription factor in the differential response observed after treatment with SIINFEKL, expression of major E2F1 targets(40) was assessed. Upregulation of most of E2F1 targets was observed in response to all treatments besides SIINFEKL peptide (Fig. 4D). The trend observed from the heatmap was validated by enrichment analysis, where *E2f1* and E2F1 targets were shown to be overrepresented in model EOT_13 (p-value: 3.1E-05).

Together, treatment with erythrocyte-targeted antigens and soluble OVA induced a similar cell-cycle program as the one described in self-reactive T cells maintained in an unresponsive state in the periphery. Surprisingly, treatment with soluble SIINFEKL peptide did not lead to such a signature, with lack of significant upregulation of *E2f1* and most of E2F1 targets.

### Nur77 (Nr4a) family of transcription factors

In the previous two sections, genes induced by erythrocyte-targeted as well as either OVA or SIINFEKL peptide were analyzed and selective regulation of the *Egr2-Nfatc1* (Fig. 4C) and *E2f1* (Fig. 4D) transcription factors in response to the different treatments was highlighted. Several genes related to peripheral tolerance were, however, solely induced in response to treatment with erythrocyte-targeted antigens, and were shown to cluster in model ET_10. Most strikingly, tolerance induction by erythrocyte-targeted antigens involved upregulation of the pro-apoptotic Nr4a family of orphan nuclear receptors members *Nr4a1* (Nur77), *Nr4a2* (*Nurr1*), and *Nr4a3* (*Nor1*) (Fig. 4E-F). This family of transcriptional factors participates in the process of apoptosis in part through interaction with the anti-apoptotic protein BCl-2 (41). While *Bcl2* gene was shown to be downregulated in response to all treatments, an increase in expression of Nr4a transcription factors was solely induced in response to erythrocyte-targeted antigens (Fig. 4E-F).

In addition to the two inhibitory receptors *Cd200r1* and *Tigit* mentioned previously, several other genes associated with the process of immune tolerance were shown to be specifically overexpressed in response to erythrocyte-targeted antigens (Fig. 4E-F). Among them, the gene encoding for the cytokine lymphotactin (Xcl1) (17), reported as being overexpressed during anergy, exhaustion and induction of regulatory T cell, was among the most highly upregulated gene in response to erythrocyte-targeted antigens compared to SIINFEKL peptide or soluble OVA. From gene ontology analysis, treatment with erythrocyte-targeted antigens was shown to induce upregulation of genes associated to IL-10 production (Fig. 2C). Interestingly, genes associated with this biological process, i.e. Cd83, Xcl1 and Tigit, showed similar gene expression features as revealed from the scatterplot representation of genes from model ET_10 (Fig. 4F).

## Discussion

Transcriptome analysis in CD8+ T cells undergoing immune tolerance in response to treatment with erythrocyte-targeted antigens, soluble OVA and soluble SIINFEKL MHC I peptide was performed in this study. While sharing common features, such as upregulation of genes associated with cell cycle and proliferation (Fig. 2B), several master transcription factors involved in the process of peripheral T cell tolerance were specifically expressed in response to the different treatments.

Relatively to in their naïve state, CD8+ T cells in mice treated with erythrocyte-targeted antigens exhibited an enhanced expression of genes encoding for TCR co-inhibitory receptors, showed impaired cytolytic gene expression and specific modulation of transcription factors involved in the TCR signaling pathway (Fig. 4). Through comparison with gene datasets characteristic of T cells undergoing activation or tolerance, treatment with erythrocyte-targeted antigens was shown to induce a tolerogenic signature in part through modulation of the transcription factors *Nr4a* family, *Egr2-Nfatc1* and *E2f1* (Fig. 4). While results obtained in this study will need to be validated at the protein level, the high consistency and similarity between previously reported gene profiles of peripherally tolerized CD8+ T cells and gene response observed after treatment with erythrocyte-targeted antigens support the idea that this strategy mimics the natural mechanisms of induction of immune tolerance.

Intravenous administration of soluble antigen OVA was shown to induce upregulation in CD8+ T cells of *GzmB* expression (Fig. 4C). Overexpression of GzmB in response to soluble OVA has been previously reported working with flow cytometry analysis (11), and would suggest that CD8+ T cells are not completely functionally impaired after intravenous administration of the soluble OVA. As no change in expression of master transcription factors driving cytotoxic response was observed, upregulation of *Gzmb* expression seemed to be rather due to the downregulation of its negative regulator BCL-6 (Fig. 4B). In addition, and compared to treatment with erythrocyte-targeted antigens, CD8+ T cells in mice treated with OVA lacked expression of several inhibitory receptors genes such as *Btla, CD200r1* and *Tigit* (Fig. 4B). Finally, as observed through comparison of our dataset with the gene set characteristic of CD8+ T cells undergoing deletional tolerance (17), CD8+ T cells in the OVA treated group did not upregulate expression of *Egr2* and *Nfatc1* transcription factors, binding partners and targets (Fig. 4), known to be involved in the process of immune tolerance.

Among the different antigen formulations tested, the transcriptional signature in response to soluble SIINFEKL peptide was the most distinct, highlighted by both the distance in the PCA (Fig. 1C) and percentage of genes clustered in model ETS_15 of our model selection analysis (Fig. 2A). Administration of a high dose of soluble peptide is known to induce T cell unresponsiveness and death (47)(48). This strategy is currently used for allergy desensitization (49) and under clinical trials for treatment of celiac disease (50). As soluble peptides can be directly loaded on MHC molecules without requirement of further processing by dendritic cells, the mechanism is very different from processing of erythrocyte-targeted antigens, thus we expect that the induced T cell response will also differ. Similarly to treatment with erythrocyte-targeted antigens, intravenous administration of soluble MHCI SIINFEKL peptide induced expression of genes associated with deletional tolerance (17) (Fig. 3). However, and as observed from comparison of our dataset with the list of genes specifically expressed in self-reactive CD8+ T cell maintained in an unresponsive state in the periphery (18), the cell-cycle program of CD8+ T cells in the SIINFEKL treated group was revealed to be very distinct from that of mice treated with OVA or erythrocyte-targeted antigens, with lack of modulation of *E2f1* transcription factor and targets (Fig. 4D). This discrepancy in cell cycle genes modulation could also reveal differences in cell-cycle kinetics, where CD8+ T cells treated with SIINFEKL peptide would proliferate more quickly after intravenous administration and enter earlier in an apoptotic phase compared to in response to full OVA or erythrocyte-targeted antigens.

Finally, genes encoding for the orphan nuclear receptors Nr4a family were shown to be solely overexpressed in response to treatment with erythrocyte-targeted antigens (Fig. 4E-F). Acting as transcription factors, NR4A receptors regulate expression of genes associated with a number of biological processes such as proliferation, metabolism, and DNA repair as well as apoptosis (51). NR4A receptors play an essential role in lymphocyte selection during central tolerance, translating the TCR signaling strength into induction of apoptosis or FOXP3 expression for differentiation into natural Tregs (46). In addition to their implication in the process of central tolerance, overexpression of three members of the NR4A receptor family has been reported in CD8+T cells undergoing deletional tolerance, suggesting a role of these transcription factors in induction of apoptosis during peripheral tolerance (17). Interestingly, neither treatment with OVA nor with SIINFEKL peptide led to the upregulation of these transcription factors, whereas administration of erythrocyte-targeted antigens led to the upregulation of all three members of the NR4A family (Fig. 4E-F).

To further understand the tolerogenic events happening in response to erythrocyte-targeted antigens, additional work would be required to determine if the induced response is dose-dependent. Indeed, in both preclinical studies in animal models as well as human clinical trials (2)(52), the dose of antigen administered has been shown to dictate the efficacy of therapy. Modification of the dose can potentially affect the amplitude of the response at the gene level. In addition, using apoptotic cells as tolerogenic carrier, others have demonstrated that the dose of the tolerogenic compound can have an impact on the “kind” of tolerance induced, from deletional tolerance/anergy to differentiation into Tregs (53). Thus, further work would be required to understand how modulation of the dose of the different formulations would impact on the gene response. The studies performed here were all at equimolar dose, at an equivalent of 10 μg of OVA antigen.

One limitation of the type of analysis performed in this study, in addition to the small number of samples per groups that may reduce the power of our data, is the lack of information regarding small populations. Indeed at the protein level, differences between treatment groups is often observed as a variation in the proportion of cells expressing certain markers, such as the inhibitory receptors CTLA4 and PD1, or the regulatory T cell marker FOXP3, rather than a change in amplitude of expression of these molecules (10, 11). Thus, single-cell analysis or flow cytometry staining would allow determination if the differences in gene expression observed in this study arise from an increased proportion of positive cells, or a global increase in gene expression.

Finally, as regulatory T cells have been shown to play a preponderant role in the maintenance of the tolerogenic state after treatment with this erythrocyte-targeted antigens (11), a similar study looking this time at gene response in CD4+ T cells could be performed to assess genes and pathways involved in the induction of FOXP3+ regulatory T cells.

In summary, this study brings novel insights into the process of induction of peripheral tolerance by erythrocyte-targeted antigens (illustrated in Fig. 4A). Involvement of the transcription factors EGR2/NFATC1, E2F1 and Nur77 family has been revealed as well as similarity in gene response compared to the natural event of induction of peripheral tolerance. In addition to bringing theoretical knowledge on the mechanisms behind induction of immune tolerance by erythrocyte-targeted antigens, this study highlights novel markers and targets that could be used to evaluate the state of tolerance induced and thus efficacy of the therapy in different situations.

## Acknowledgments

We thank J-P. Gaudry, K. Brünggel, Dr. E. Simeoni, Dr. Z. Julier and Dr. A. de Titta for their help and useful discussions.

